## Supplemental Figures for Figures 1 and 2 for "A dysmorphic mouse model reveals developmental interactions of chondrocranium and dermatocranium"

Figure 1 – Figure supplement 1

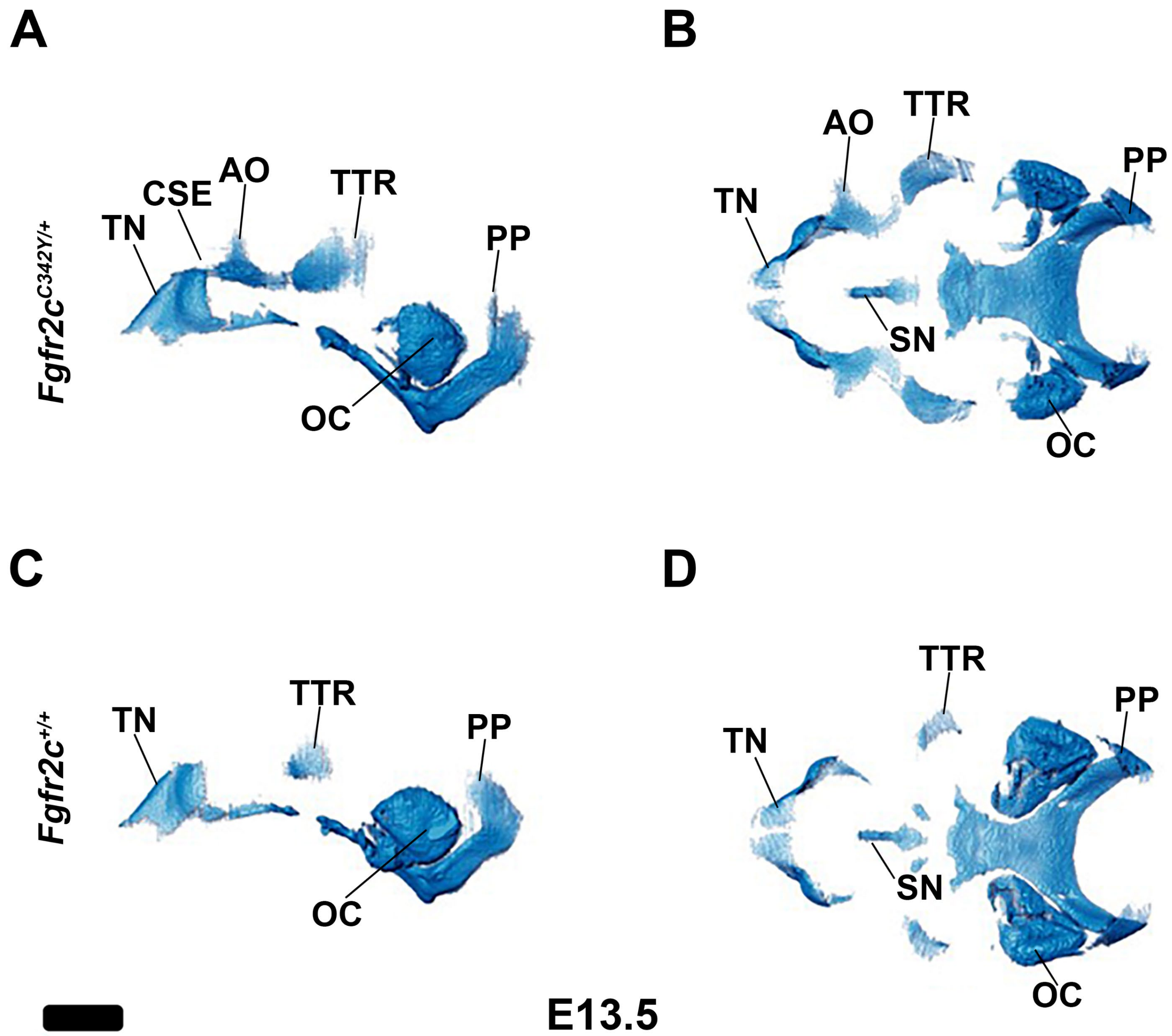

Figure 1 – Figure supplement 2

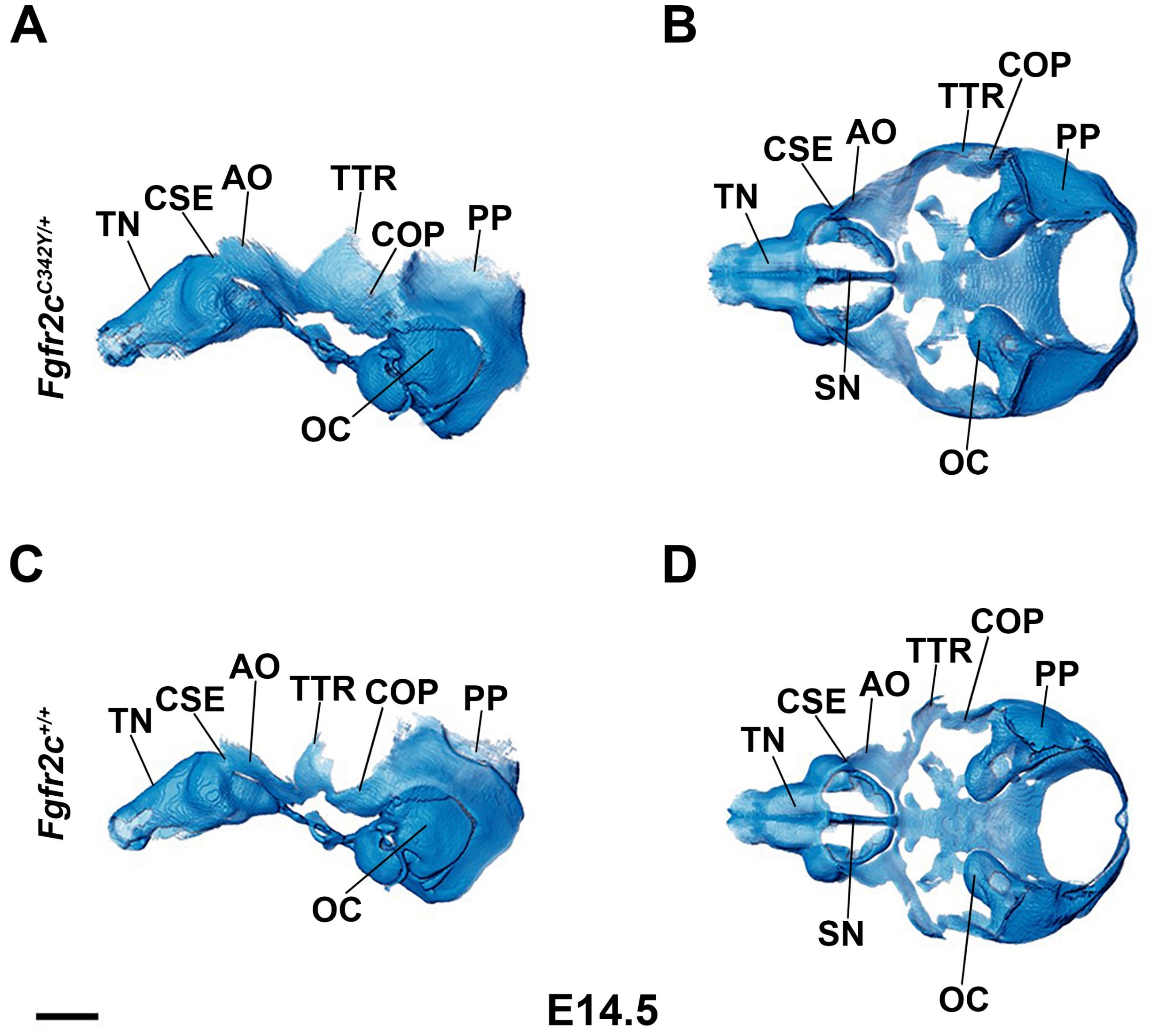

Figure 1 – Figure supplement 3

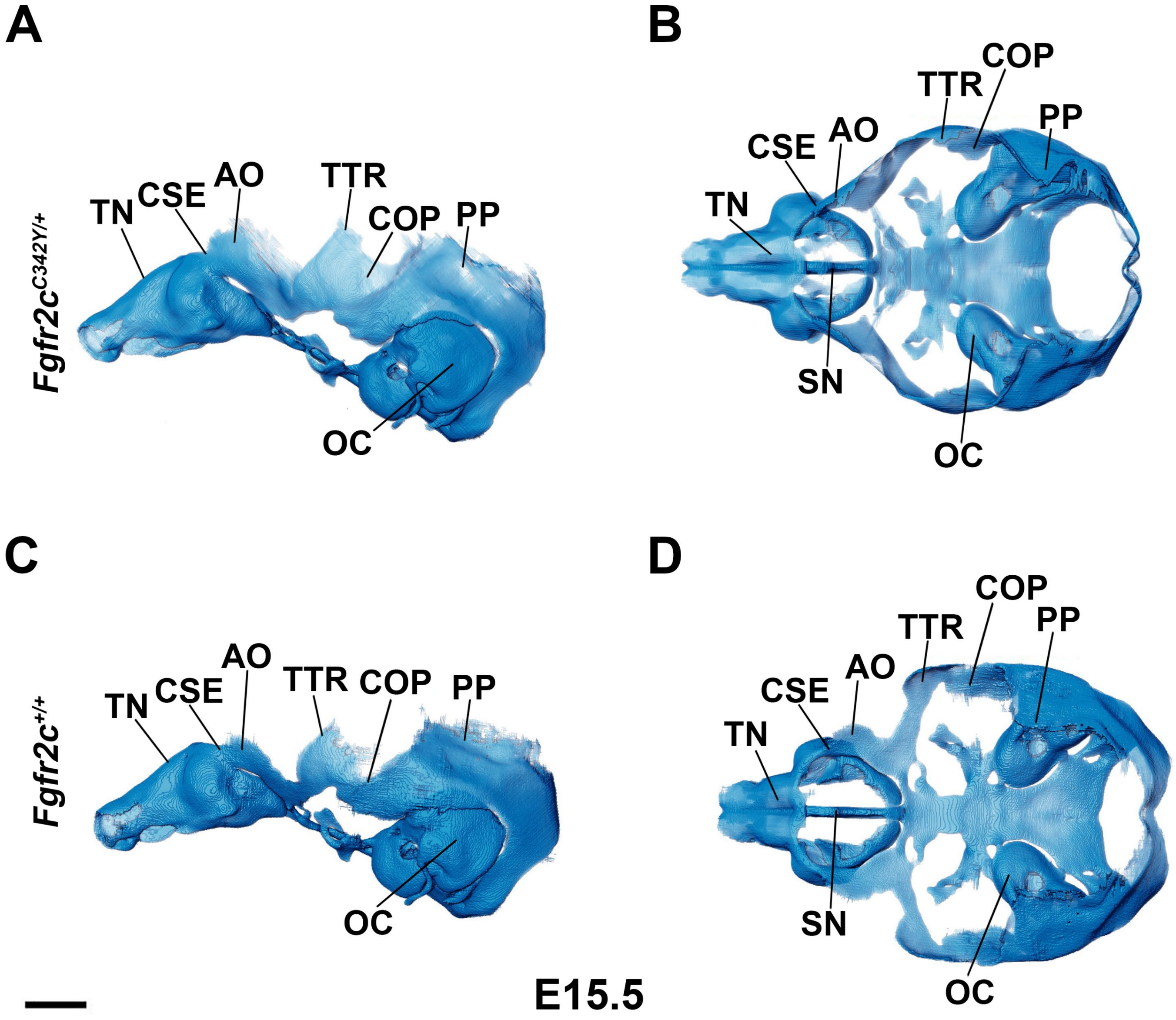

Figure 1 – Figure supplement 4

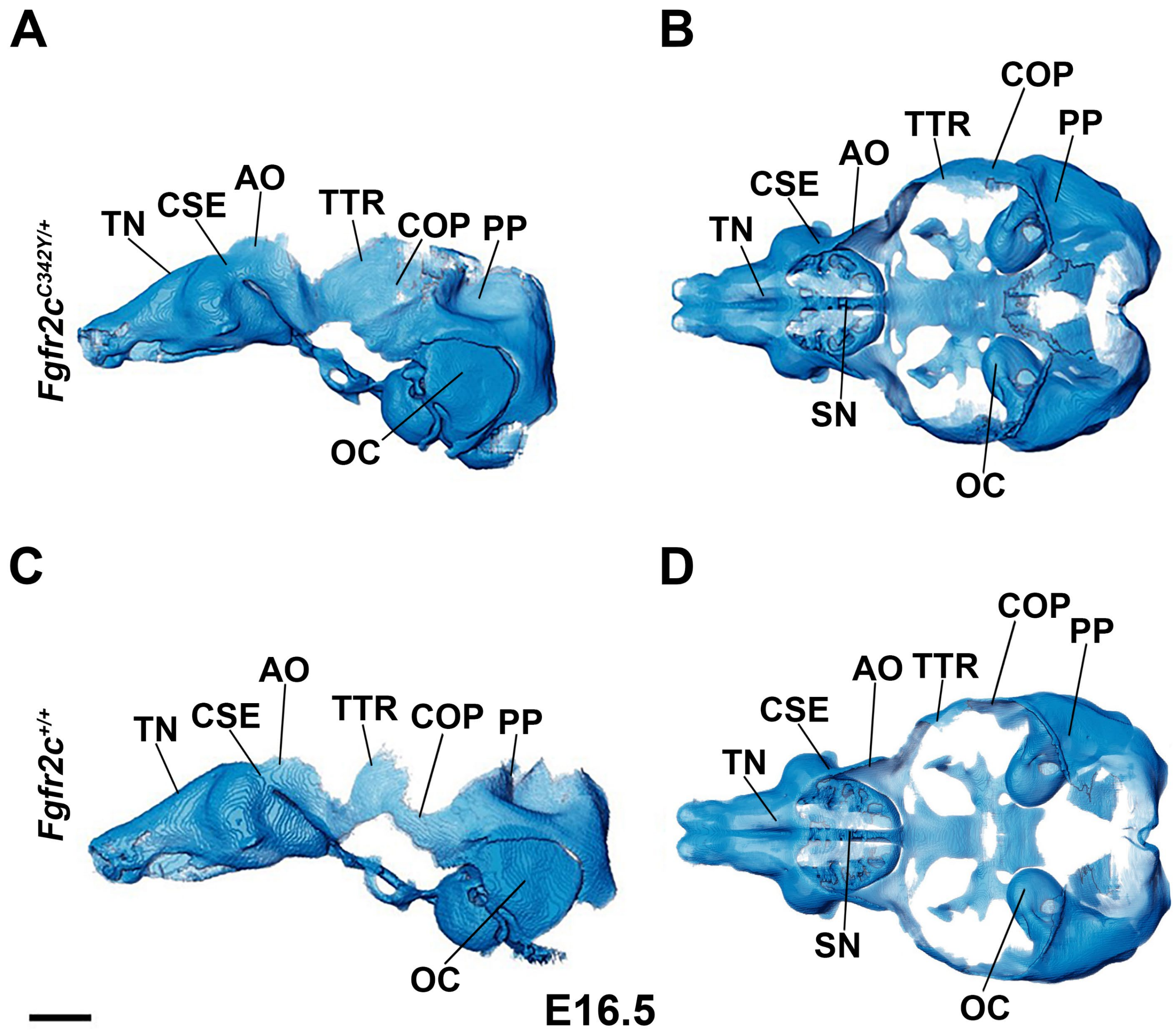

Figure 1 – Figure supplement 5

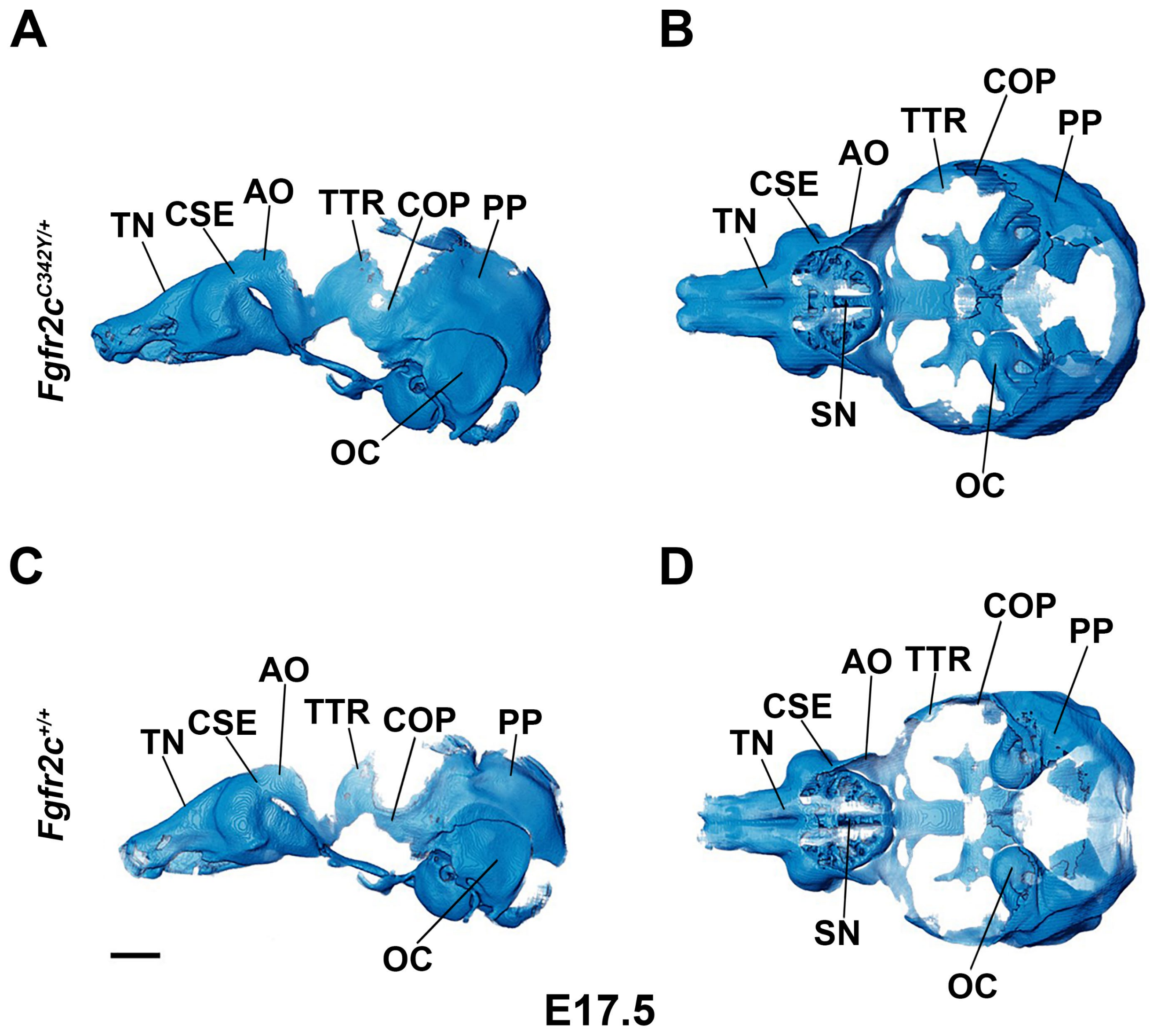

Figure 2 – Figure supplement 1

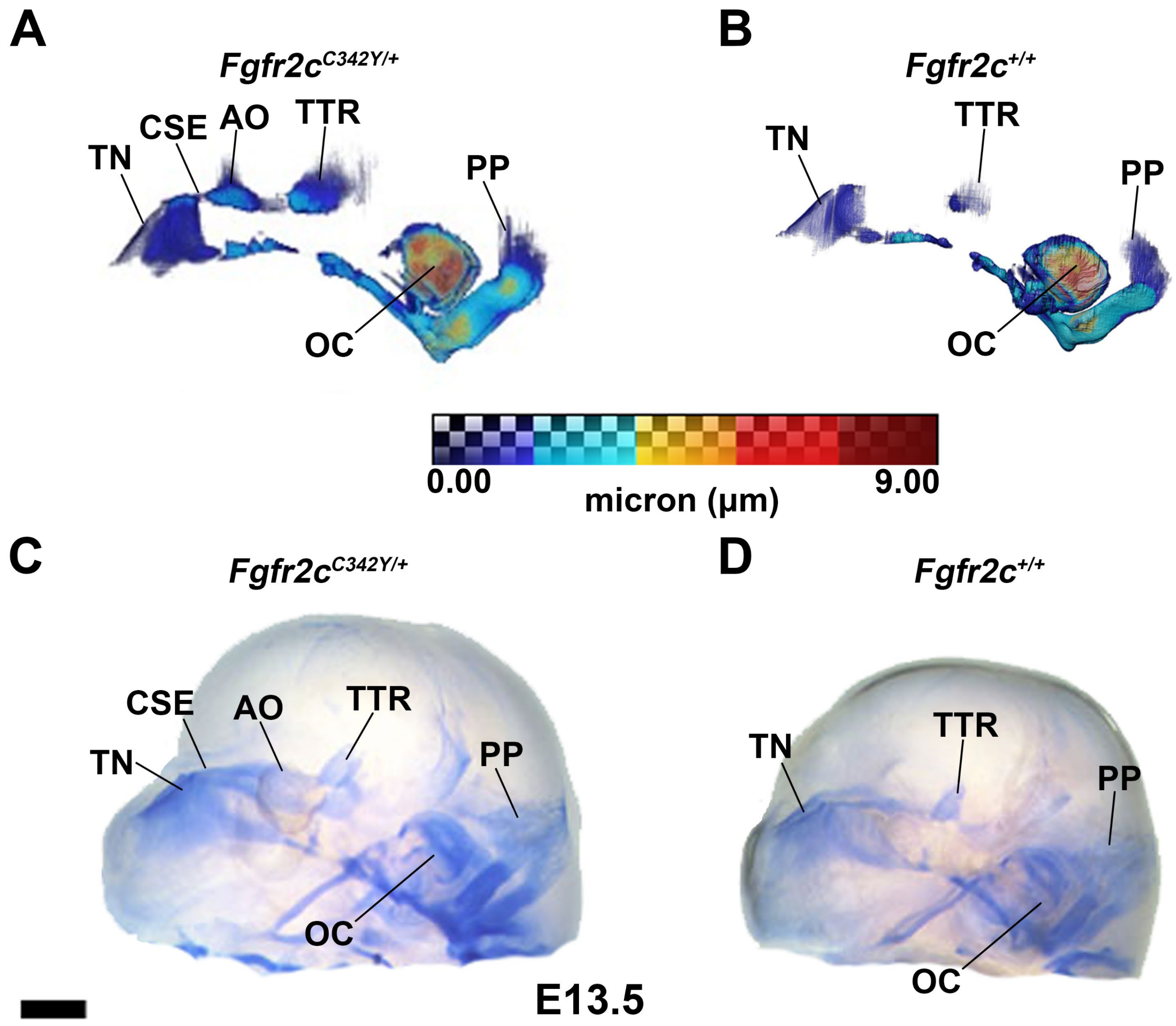

Figure 2 – Figure supplement 2

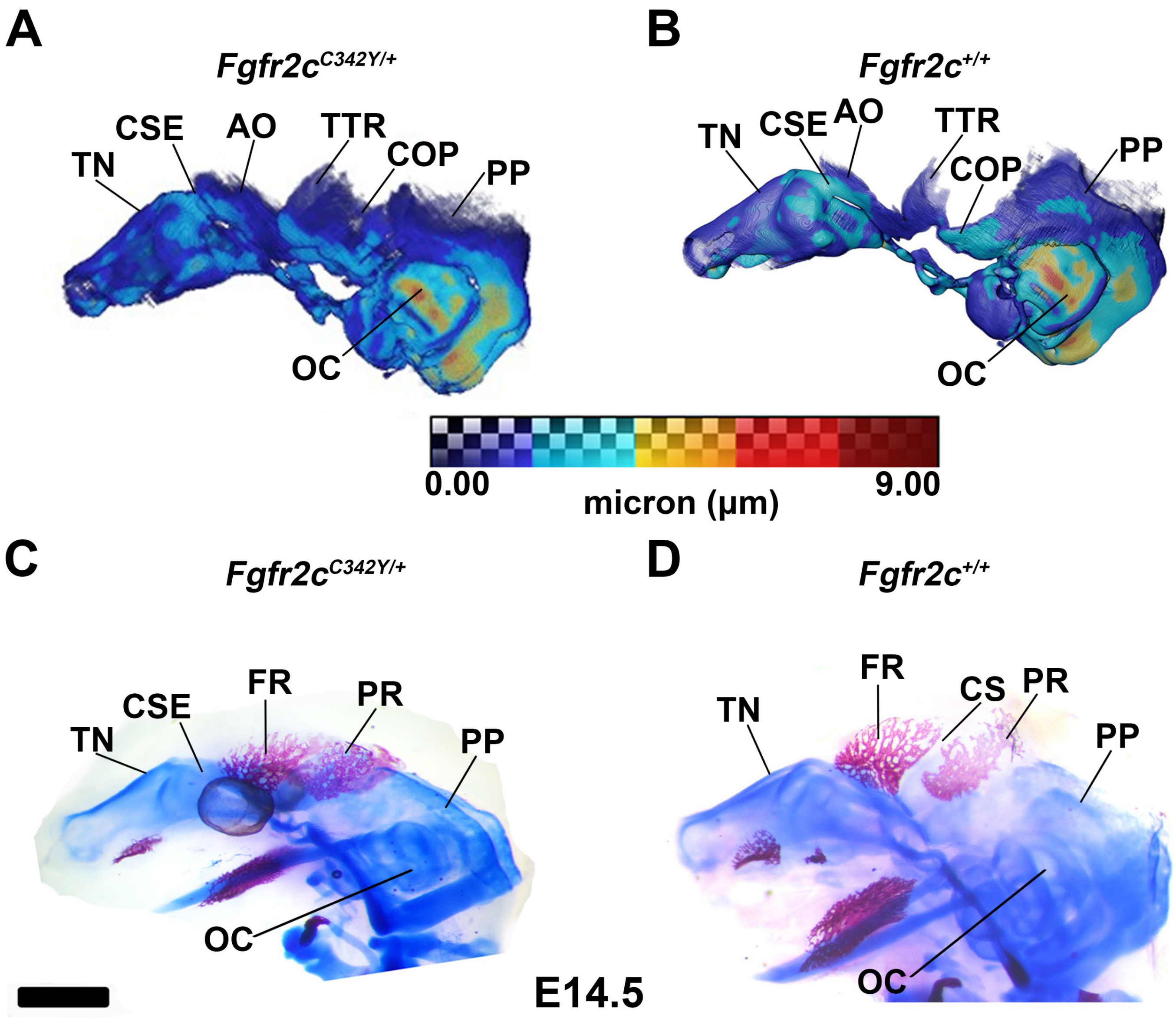

Figure 2 – Figure supplement 3

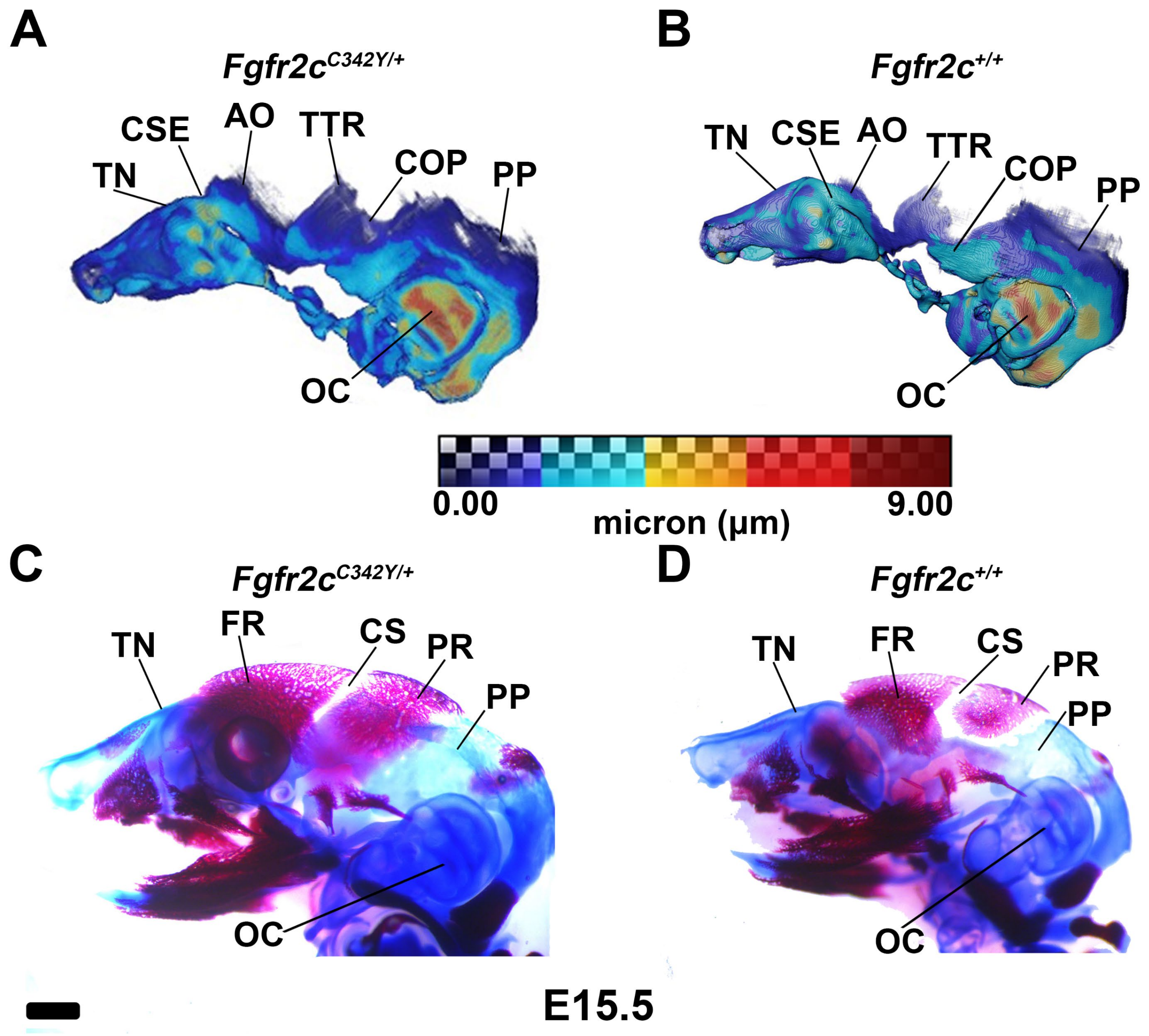

Figure 2 – Figure supplement 4

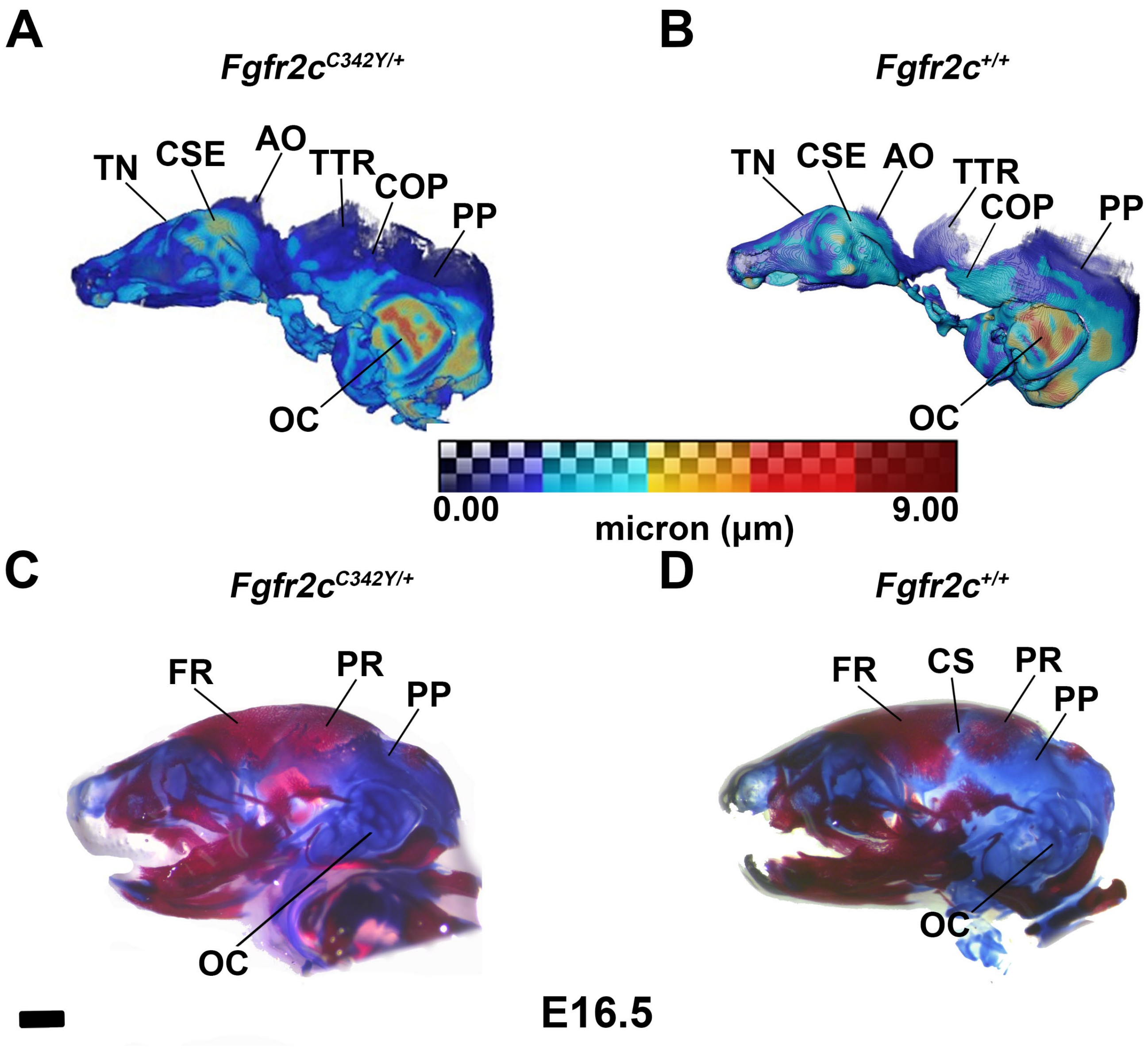

Figure 2 – Figure supplement 5

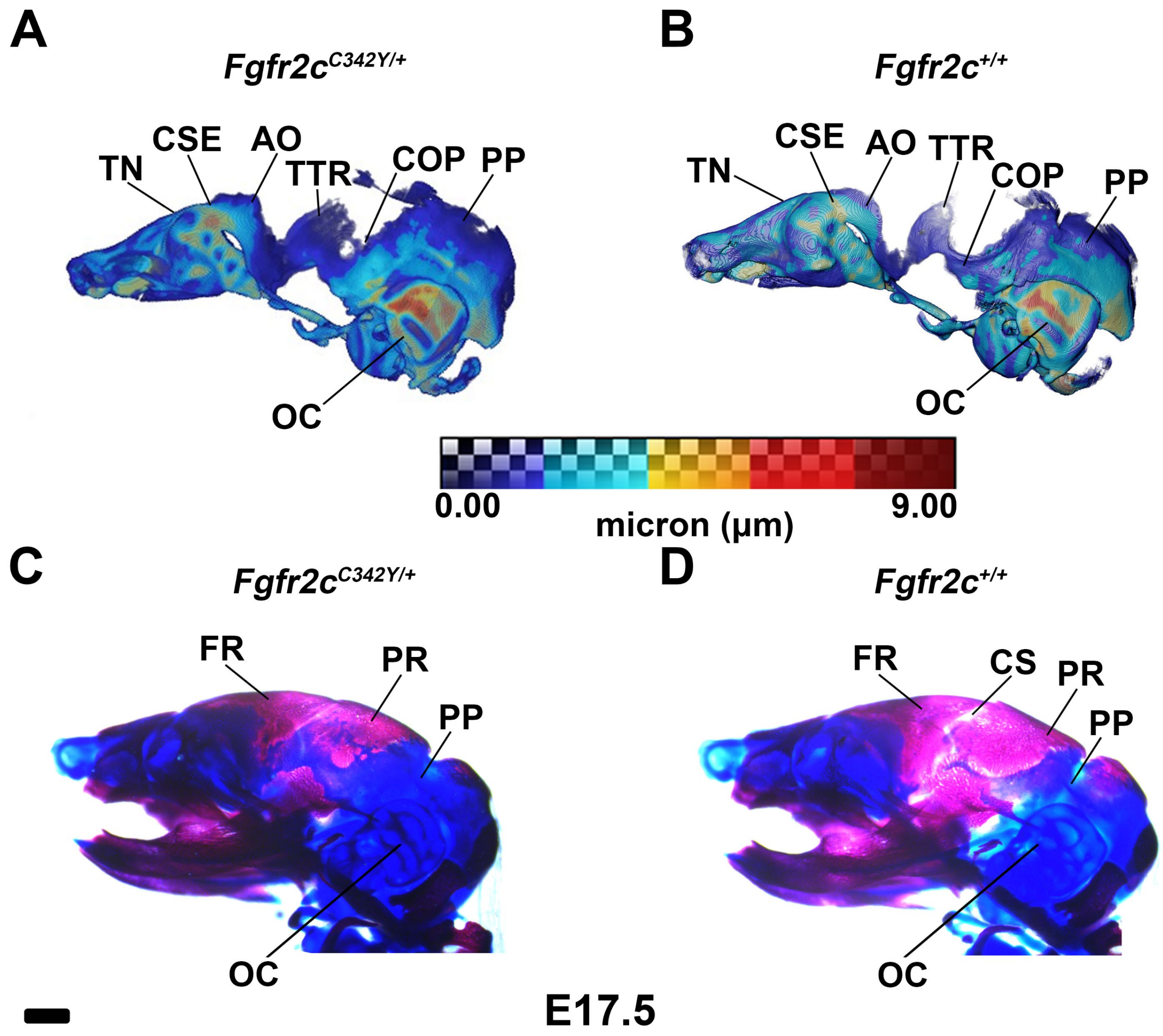
